# Multilayer Network Modelling of the Human Reading System

**DOI:** 10.64898/2026.01.19.700433

**Authors:** Vicky He, Mangor Pedersen, David N. Vaughan, Heath R. Pardoe, Jodie E. Chapman, Graeme D. Jackson, David F. Abbott, Chris Tailby

**Affiliations:** The Florey Institute of Neuroscience and Mental Health, Heidelberg, Victoria, Australia; Florey Department of Neuroscience and Mental Health, The University of Melbourne, Parkville, Victoria, Australia; Department of Psychology and Neuroscience, Auckland University of Technology, Auckland, New Zealand; Department of Neurology, Austin Health, Heidelberg, Victoria, Australia; Department of Medicine-Austin Health, The University of Melbourne, Heidelberg, Victoria, Australia; Department of Clinical Neuropsychology, Austin Health, Heidelberg, Victoria, Australia

## Abstract

Reading is supported by rapid and flexible coordination of neural activity across distributed brain regions. We have previously shown that left fusiform gyrus (FusG) provides a bridge flexibly linking the visual form analysis required for reading with the language system. Here, we investigate the dynamic organisation of an extending reading network encompassing classical perisylvian language areas and FusG. We do so by applying multilayer network modelling to language fMRI data acquired through the Australian Epilepsy Project, using a paradigm that contrasts reading with visuospatial judgements. The dataset included 201 participants with left dominant language, both with and without seizure disorders. We hypothesised that the relative strength of dynamic interactions within this extended language network is associated with reading ability. Time resolved functional connectivity was estimated using a sliding window Pearson’s correlation approach, and the resulting connectivity matrices were entered into a multilayer community detection algorithm to quantify spatiotemporal community structure within the reading network. We concentrate our analyses on *allegiance*, the probability that a pair of regions is assigned to the same community over time. Our results show that community structure within the reading network is characterised by a preference for within hemisphere assignment over cross hemisphere assignment, as well as higher nodal allegiance among left language regions compared with their right hemisphere homologues. As anticipated, within versus between network allegiance followed a similar gradient in both language and attention networks: lowest between left language and right attentional regions, intermediate between the left FusG and each respective network, and highest within-network (left language or right attention). Importantly, as hypothesised, reading ability was associated with FusG-inferior frontal gyrus (IFG) interactions: higher left FusG-left IFG allegiance correlated with better reading performance, whereas increased right FusG-left IFG allegiance correlated with poorer reading. These findings highlight hemispheric asymmetries in the dynamic organisation of the reading system and provide novel evidence linking individual differences in reading ability to network level dynamics. Our findings align with a developmental literature suggesting that as reading proficiency improves, there is a shift from bilateral to unilateral left occipitotemporal engagement.

## 1 Introduction

The human reading system, like other cognitive systems, is inherently dynamic. During reading, visual text is processed along the visual form pathway through left fusiform gyrus (FusG), where the Visual Word Form Area is found (Cohen et al., 2002; Baker et al., 2007) and relayed into perisylvian language regions for linguistic processing (Price, 2012; Chen et al., 2019; He et al., 2025a). Processing of non-text visual inputs still recruits visual form areas, though in the absence of text or grapheme-based inputs the need for coupling to language areas is reduced if not eliminated. Thus, the integration of visual form analysis regions and classical language regions is likely to vary over time as a function of the nature of visual inputs and the behavioural responses required to those inputs.

Multilayer network modelling (MLNM) is a recently developed approach for quantifying the spatiotemporal organisation of brain networks (Bassett et al., 2011; Bassett et al., 2013a; Mucha et al., 2010). It is based on the idea that brain nodes are dynamically organised into modules to support specific functions, where a node represents a brain region and a module is defined as a group of nodes that preferentially connect to each other during a particular temporal window (Bassett et al., 2011). This method relies on a community detection algorithm that maximises a multilayer modularity quality function to identify a spatiotemporal layout (i.e., nodes are interconnected across both space and time) that best partitions brain nodes into time-varying modules (Bassett et al., 2011; Bassett et al., 2013a; Kelkar & Medaglia, 2018). One commonly derived summary measure from this approach is *allegiance*, which is defined as the probability that a pair of nodes is assigned to the same module across time. The collection of these pairwise values across the network constitutes the module allegiance matrix.

Using measures such as module allegiance, as well as related statistics describing the stability and flexibility of community assignments, prior research has identified links between dynamic modular network organisation and cognitive function. Examples include evidence of a core-periphery structure in the language system during reading (Chai et al., 2016), the emergence of modular structure over the course of learning (Bassett et al., 2013b; Bassett et al., 2015), as well as predictive relationships between network switching rates and task performance (Pedersen et al., 2018b).

Previous work on the language systems has tended to focus primarily on core perisylvian regions, proximal to Broca and Wernicke areas (e.g., Chai et al., 2016; He et al., 2018; Ke et al., 2023). Comparatively less work has examined how left FusG, commonly regarded as a sensory processing region rather than part of language per se, integrates with the classical speech-based language system. However, FusG plays a critical role in flexibly linking visual input with language and attentional systems (Chen et al., 2019; He et al., 2025a), as well as in the development of reading proficiency. The visual processing that supports reading is thought to develop bilaterally in the ventral occipitotemporal pathway (where FusG is located), and as people become more proficient in reading, this system becomes increasingly left lateralised (Tailby et al., 2014; Shaywitz et al., 2002). MLNM provides a framework for characterising how the left FusG dynamically interacts with perisylvian language regions.

In this work, we use MLNM to test three specific hypotheses involving the left FusG. First, in individuals with left-dominant language, we expect left language regions to show higher allegiance compared with their right hemisphere homologues (H1). To be consistent with past research (He et al., 2018), intrahemispheric allegiance, defined as the tendency for regions within the same hemisphere (e.g., left language regions) to be assigned to the same community over time, is referred to as *recruitment*. Similarly, interhemispheric community assignment (e.g., left language-right language homologues) is referred to as *integration*. This hypothesis therefore predicts greater recruitment among left language regions and relatively lower integration across hemispheres. Relatedly, we also expect stronger left language-left language recruitment compared with right language homologue-right language homologue recruitment. Second, in previous work using psychophysiological interaction (PPI) analysis, we showed up-regulation of information flow between the left FusG and left language regions during reading, as well as between the left FusG and right visuospatial attentional regions during pattern matching (He et al., 2025a). We therefore hypothesised a gradient of allegiance among left language, right attentional, and left FusG regions: lowest between left language and right attentional regions, intermediate between left FusG and left language/right attentional regions, and highest within network (left language or right attentional; H2). Finally, visual processing for reading develops bilaterally in the ventral occipitotemporal pathway and becomes increasingly left lateralised with greater reading proficiency (Tailby et al., 2014; Shaywitz et al., 2002). Accordingly, we expect a positive relationship between reading ability and the probability of left FusG-left language recruitment, and an inverse relationship between reading ability and the probability of right FusG-left language integration (H3).

## 2 Methods

### 2.1 Participants

Participant characteristics are described in He et al. (2025a, 2025b). The present study uses the same participant sample as reported in those papers. We note that we are leveraging data from an existing project, the Australian Epilepsy Project (AEP), to test hypotheses regarding the human reading system. In brief, the study sample comprised adult participants recruited through the AEP. The analysis cohort included 94 participants with seizure disorders (median age 33 years, interquartile range = 19; 47 males), and 107 healthy controls (median age 43 years, interquartile range = 22; 38 males). Ethics approval for the study was obtained from the Austin Health Human Research Ethics Committee (HREC/60011/Austin-2019; HREC/68372/Austin-2022).

All participants had a functional level of English and left-lateralised language. Language lateralisation was determined based on methods described in Abbott et al. (2010). As part of the AEP neuropsychological assessment completed by the present sample, participants were administered the Test of Premorbid Functioning (ToPF; Wechsler, 2011), which assesses the pronunciation of irregular words. This measure was used as an estimate of reading ability, and throughout this manuscript we refer to ToPF scores as reading scores (normative mean *± SD* = 100 *±* 15). Reading scores were available for 89 of the 94 participants with seizures and 94 of the 107 healthy controls. All participants have at least a basic level of reading ability (Table 1), making their inclusion valid for testing reading-related hypotheses. Combining the seizure and healthy volunteer cohorts ensures a broad range of reading abilities, making the sample valuable for exploring the relationship between reading performance and network metrics. In all modelling, we include a group factor (seizure disorders, healthy controls), though our principal interest is in reading related network dynamics. Relevant sample characteristics are summarised in Table 1. Further details of the cohort can be found in Table 1 in He et al. (2025a).

**Table 1:** Participant characteristics.^a^.

| Participant characteristics | Seizure<br>($N = 94$) | Control<br>($N = 107$) | Test-statistics | p-value |
| --- | --- | --- | --- | --- |
| Age (years) | 33 (19.5) | 43 (22.0) | $W = 3395$ | $< 0.001$ |
| Males | 47 (50.0) | 38 (35.5) | $\chi^2 = 3.73$ | 0.053 |
| Reading scores | 102 (22.0) | 111 (12.8) | $W = 2822$ | $< 0.001$ |
| Education (years) | 15 (3.0) | 15 (3.0) | $W = 3262$ | $< 0.001$ |
| IQ | 101 (19.0) | 110 (17.5) | $W = 2694$ | $< 0.001$ |

<sup>a</sup> Age, reading scores, education in years, and IQ are given in median (interquartile range, IQR); sex (number of males) is given as count (%).

### 2.2 MRI data acquisition and preprocessing

MRI details are described in He et al. (2025a). In brief, we collected both T1-weighted and Multi-Band Multi-Echo (MBME) fMRI images for all participants in a 3T Siemens PrismaFit MRI scanner, with the following parameters: TR = 0.9 s, multi-band factor = 4, and voxel size 3 × 3 × 3 *mm*. Image preprocessing included Marchenko-Pastur PCA (MPPCA) denoising (Veraart et al., 2016) implemented in MRtrix3 v3.0.4 (Tournier et al., 2019) to remove thermal noise (Ades-Aron et al., 2021; Mosso et al., 2022). This was followed by preprocessing using fMRIPrep 21.0.2 (Esteban et al., 2019, 2020). An 8 mm full width at half maximum Gaussian kernel was used to smooth all functional images.

### 2.3 fMRI paradigm

Participants completed a block design fMRI task contrasting pseudoword rhyming against pattern matching. In the rhyming blocks, participants were asked to decide whether visually presented pairs of pseudowords rhyme or not (e.g. “blint” vs “glat”). In the pattern matching blocks, participants were asked to judge whether pairs of line patterns were identical or not (e.g., “//*\\*” vs. “*\\\*/”). There was no separate resting baseline.

Each task block consisted of four pairs of visual stimuli, and each pair was displayed for 4.5 seconds. There were 20 rhyming and 20 pattern pairs in total. The fMRI experiment lasted 180 seconds (200 TRs; TR = 0.9 s). The first 10 volumes were discarded from the subsequent analysis to allow participants to settle. The total number of volumes used for subsequent analysis is 190.

### 2.4 Network construction

#### 2.4.1 Parcellation and time series extraction

Brain nodes were defined using a whole brain parcellation based on the Brainnetome Atlas (BNA), which comprises 246 nodes delineated on the basis of functional connectivity (Fig. 1a; Fan et al., 2016). Previous research has suggested that this atlas is suitable for network-based analysis (Breedt et al., 2022). As BNA is a probabilistic atlas, parcels were thresholded at 50%.

**Figure 1:**
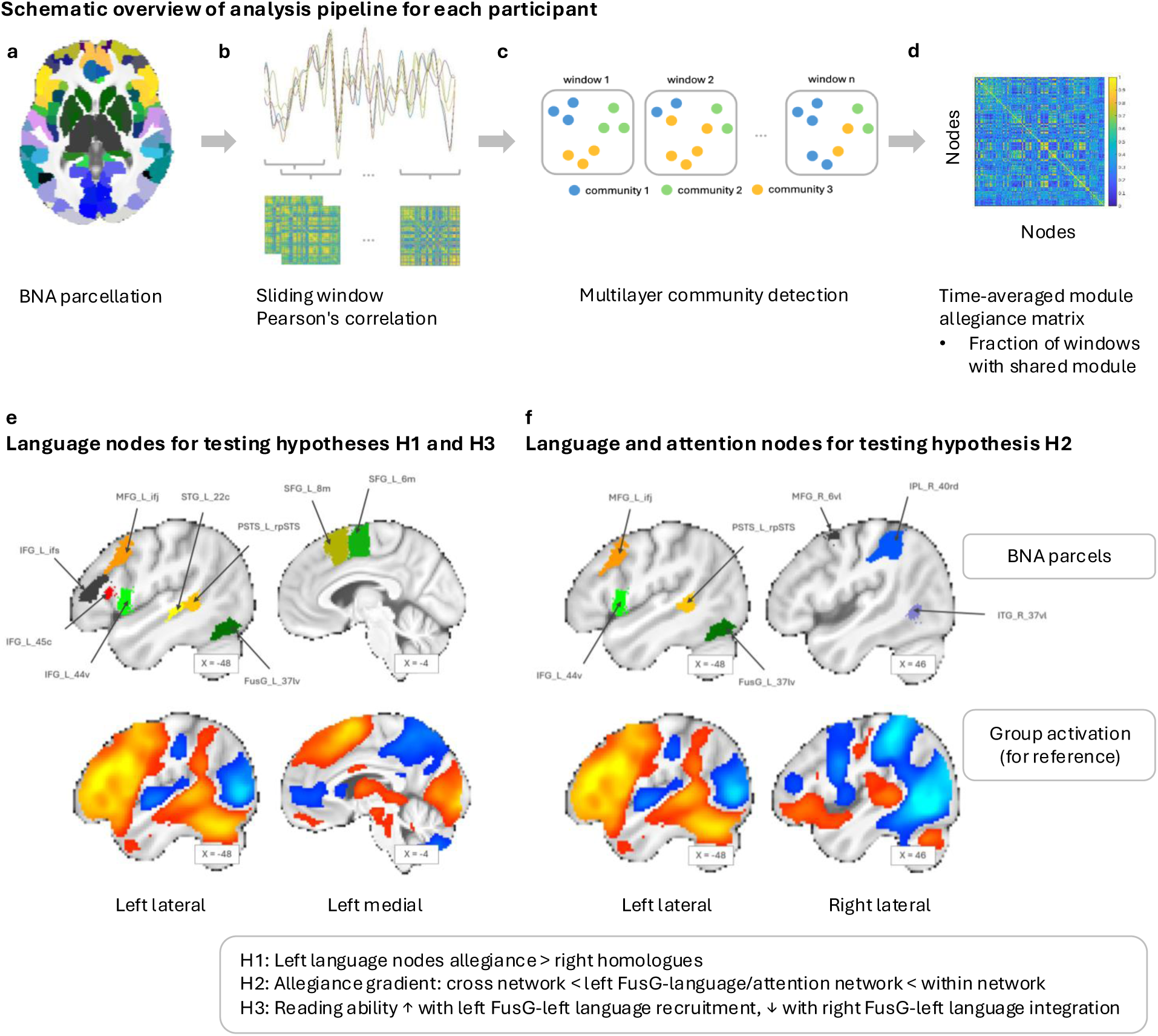
Schematic overview of the analysis pipeline for each participant. a. Whole-brain parcellation using BNA, resulting in 246 brain nodes. b. Sliding-window Pearson’s correlation approach (top) used to estimate adjacency matrices (bottom). c. Illustration of multilayer community detection algorithm output, where each community represents a *module*. d. Time-averaged module allegiance matrix based on community assignments, with each cell representing the fraction of windows that the corresponding row and column regions were assigned to the same community. e. Anatomical locations of the selected language regions in the left hemisphere used to test hypotheses H1 and H3. f. Anatomical locations of the selected language (left hemisphere) and attention regions (right hemisphere) used to test hypothesis H2. Region definitions are provided in Table 2.

We extracted a time series for each of the 246 nodes using SPM12 (revision 7771; Friston et al., 2007). First, we performed a task activation analysis. A general linear model (GLM) contrasting pseudoword rhyming blocks (coded as 1) with pattern matching blocks (coded as 0) was fitted. The GLM also included the following confound regressors: 24 head motion parameters (Friston et al., 1996), the first eigenvariates of white matter and cerebrospinal fluid signals, and a constant term. A 128 s high-pass filter and prewhitening with a FAST model were applied. The FAST model is recommended for TRs shorter than 1 second (Corbin et al., 2018; Olszowy et al., 2019). We then regressed out the task effect along with all confound variables (Braun et al., 2015; Cao et al., 2014). From the residuals, the first eigenvariate was calculated across all voxels within each parcel, excluding voxels outside each subject’s brain mask. The resulting time series was further band-pass filtered between 0.01 and 0.1 Hz (Pedersen et al., 2018b; Ke et al., 2023) using MATLAB’s bandpass function, as low-frequency components below 0.1 Hz in fMRI data are thought to reflect functional connectivity (Sun et al., 2004; Cordes et al., 2001).

#### 2.4.2 Time-resolved community structure

We applied a sliding window Pearson’s correlation approach to estimate time varying functional connectivity between each pair of nodes, which served as inputs to the multilayer community detection algorithm (Fig. 1b). A window length of 40 TRs (36 seconds) was used for the primary analysis (He et al., 2018), as previous research has shown that windows of 30-60 seconds yield robust functional connectivity estimates (Hutchison et al., 2013; Shirer et al., 2012; Jones et al., 2012), and this value is commonly used in past research (e.g., He et al., 2018). Longer windows tend to produce more reliable results but are less sensitive to fast-scale changes in dynamic connectivity. On the other hand, shorter windows are more sensitive to transient changes, but also more susceptible to high frequency noise (Leonardi & Van De Ville, 2015; Zalesky & Breakspear, 2015). We therefore performed sensitivity analyses with shorter (20 TRs, 18 s) and longer (60 TRs, 54 s) windows.

With the applied band-pass filter, a 36 second window captures frequencies between 0.027 and 0.1 Hz. The window step was set to one TR, resulting in a total of 150 overlapping windows. Within each window, Pearson’s correlation coefficients were computed between all pairs of brain nodes after applying a Hamming window to the time series, yielding a series of time-resolved correlation matrices, hereafter referred to as adjacency matrices (Pedersen et al., 2018b). These adjacency matrices were then used as inputs to a multilayer community detection algorithm (Mucha et al., 2010). Following past research (Pedersen et al., 2018b; Suo et al., 2023), all negative correlations were set to zero to ensure that each adjacency matrix contained only non-negative entries. The multilayer community detection algorithm requires two additional parameters: the intralayer resolution parameter *γ*, which controls the size of the modules, and the interlayer coupling parameter *ω*, which controls the coupling strength between neighbouring windows (Mucha et al., 2010; Yang et al., 2021). Measures related to module allegiance were reported to be highly correlated across different values of *γ* and *ω* (Gu et al., 2022; Yang et al., 2021). We therefore chose [*γ* = 1*, ω* = 1] for consistency with most prior studies (Pedersen et al., 2018b; Ke et al., 2023; Gu et al., 2022; Suo et al., 2023; Liu et al., 2022).

The algorithm produced a time-resolved community assignment for each node in the network (Fig. 1c). Because of the stochastic nature of the community detection algorithm, the procedure was repeated 100 times (Chai et al., 2016). From these assignments, we computed a module allegiance matrix for each participant (Bassett et al., 2015; Chai et al., 2016; Ke et al., 2023), in which each cell represents the probability that the corresponding row and column regions were assigned to the same community, averaged across all windows and all iterations (Fig. 1d). By definition, this matrix is symmetrical.

The module allegiance matrix is calculated across 246 BNA nodes to obtain a reliable modular decomposition, but we restricted our hypothesis-driven analysis to nodes implicated in pseudoword rhyming and visuospatial pattern matching based on the group activation analysis described in He et al. (2025a). These regions were selected based on their overlap with the group-level language activation (and deactivation) map. The selected regions are listed in Table 2 and Figure 1e,f.

**Table 2:**
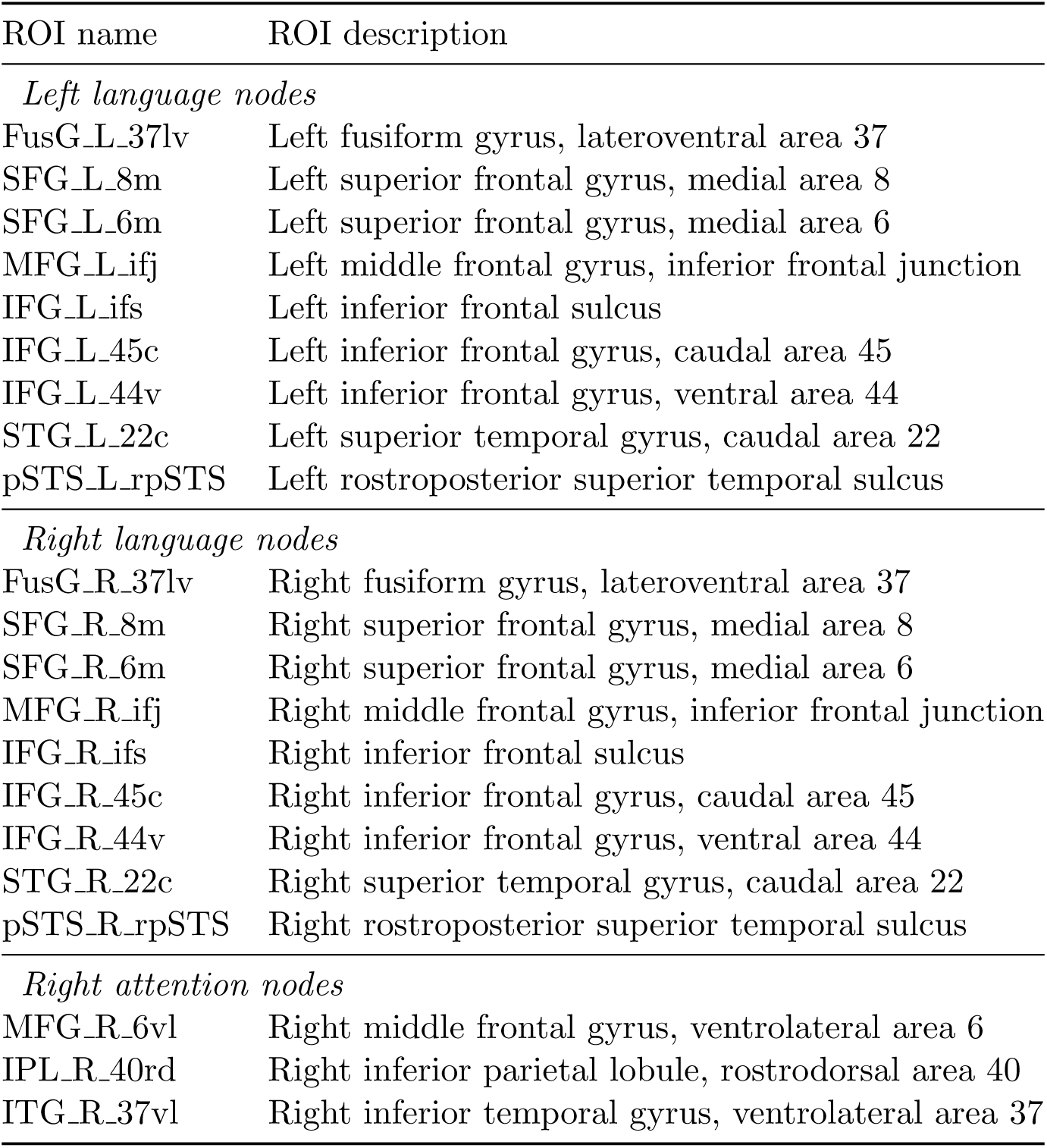
Regions of interest (ROIs).

#### 2.4.3 Lateralisation of reading modules

To examine hemispheric differences in reading network organisation (H1), we compared intrahemispheric and interhemispheric connectivity within the reading system. Specifically, to test whether the probability of left language-left language recruitment exceeded that of left language-right language integration, we extracted the subset of the module allegiance matrix corresponding to left language-left language region pairs and the subset corresponding to left language-right language region pairs (see also Fig. 2a). We then computed the element-wise difference for each participant. Inspection of two-sample t-tests comparing patients and controls indicated no significant difference in their module allegiance matrices, so they were combined into a single group for this analysis. A paired t-test was then performed on each off-diagonal matrix element, with Bonferroni correction applied (*α* = 0.05*/*72, two-sided). The diagonal was excluded because intrahemispheric elements on the diagonal are always one. Similarly, we compared left languague-left language integration against right language holomgue-right language homologue integration. Due to the symmetric nature of the module allegiance matrix, Bonferroni correction was applied with *α* = 0.05*/*36 (two-sided).

**Figure 2:**
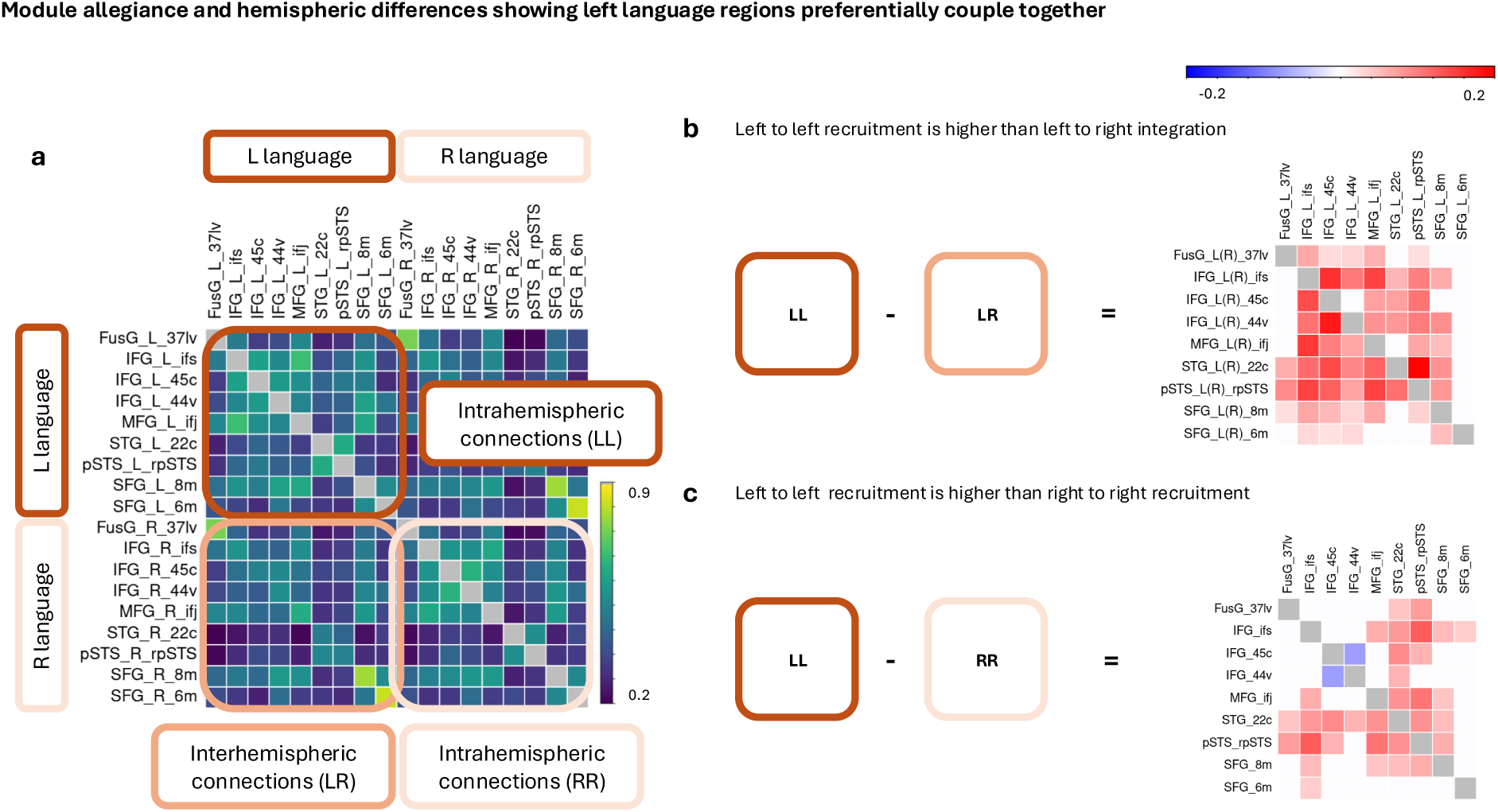
Module allegiance and hemispheric differences in language network (testing H1). a. Group-averaged module allegiance matrix. Each cell represents the probability that the corresponding row and column regions are assigned to the same community. Green box contains intrahemispheric connections with the left language regions, yellow box contains intrahemispheric connections with the right language regions, and purple box contains interhemispheric connections. Diagonals all have value one and are shaded grey. b. Differences in probability between left language-left language recruitment and left language-right language integration. White cells denote non-significant results. Cells on main diagonal are shaded white, as they were not included in the analysis. c. Differences in probability between left language-left language recruitment and right language-right language recruitment. This matrix is symmetric. White cells denote non-significant results. Cells on main diagonal are shaded white, as they were not included in the analysis.

#### 2.4.4 Quantifying left FusG coupling across language and attention networks

To examine whether the left FusG flexibly couples with left language and right attentional regions (H2), we quantified three classes of node pairs: (i) left language-right attention pairs, (ii) left FusG-network pairs (i.e., left FusG-left language and left FusG-right attention), and (iii) within-network pairs (i.e., left language-left language and right attention-right attention).

Because this analysis involved cross-network comparisons, we aimed to ensure that the included nodes were not in close proximity (e.g. adjacent BNA ROIs, which might inflate estimates of within network connectivity). We therefore selected three spatially distinct language nodes: the left MFG, IFG, and pSTS; and three spatially distinct attention nodes: the right MFG, IPL, and ITG (see Fig. 1f for node locations). For each individual, we extracted the average allegiance for each of the three node pairs defined above. We then performed paired *t* -tests separately for the language and attentional networks to compare (i) versus (ii), and (ii) versus (iii). This resulted in four paired *t* -tests in total, hence the results were Bonferroni corrected to *α* = 0.05*/*4 (two-sided).

#### 2.4.5 Recruitment and integration of left and right FusG into the left perisylvian language system and their relation to reading ability

To examine whether reading performance is related to the recruitment and integration of the left and right FusG into the left perisylvian language system (H3), we extracted allegiance values between the left FusG and other left-hemisphere language regions, as well as values between the right FusG and left-hemisphere language regions (see also Figure 4a), for each participant. These values were then entered as predictors in a linear regression model with reading score as the outcome. Age, sex, and group (patient vs. control) were included as covariates of no interest. To assess whether these relationships differed between patients and controls, we fitted an additional model including interaction terms between allegiance values and group.

## 3 Results

### 3.1 Left language regions exhibit preferential intrahemispheric recruitment over interhemispheric integration

Figure 1e shows the group-level activation map for the pseudoword rhyming task, revealing clear activation clusters in perisylvian language regions that overlap with the selected nodes in the BNA parcellation. Figure 2a shows the group-averaged module allegiance matrix. Figure 2b shows differences in allegiance between left language-left language recruitment and left language-right language integration.

The results largely support the hypothesis that left language regions are more likely to recruit within their own community than to integrate with right hemisphere homologues. Most differences favour intrahemispheric over interhemispheric connections and are statistically significant. The left SFG medial area 6 was the only region that showed no clear preference for either intrahemispheric or interhemispheric coupling. We also observed that the left FusG preferentially recruits left temporal regions but not left frontal regions, as shown in the first column in Fig. 2b. Furthermore, examining FusG connections specifically (first row in Fig. 2b), almost all left-hemisphere language regions preferentially form community with the left FusG rather than integrate with the right FusG.

The exceptions were the two superior frontal regions and the superior temporal gyrus (STG) caudal area 22, which showed no clear preference.

The majority of left language intrahemispheric recruitment also exceeded right language homologue intrahemispheric recruitment (Fig. 2c), except for the recruitment between the two IFG nodes (ventral area 44 and caudal area 45), where the allegiance among the right homologues was higher than that among the corresponding left-hemisphere regions.

### 3.2 Hemispheric gradient in left FusG coupling

The mean allegiance between left language and right attentional regions was 0.40 (*SD* = 0.09); between the left FusG and left language regions was 0.42 (*SD* = 0.12); and among left language regions was 0.47 (*SD* = 0.12). The mean allegiance between the left FusG and right attentional regions was 0.58 (*SD* = 0.14), and among right attentional regions was 0.62 (*SD* = 0.14). The observed patterns support the hypothesised gradient of allegiance, except for the comparison between left language-right attention and left FusG-left language allegiance (*p* = 0.02 *>* 0.0125; Fig. 3).

**Figure 3:**
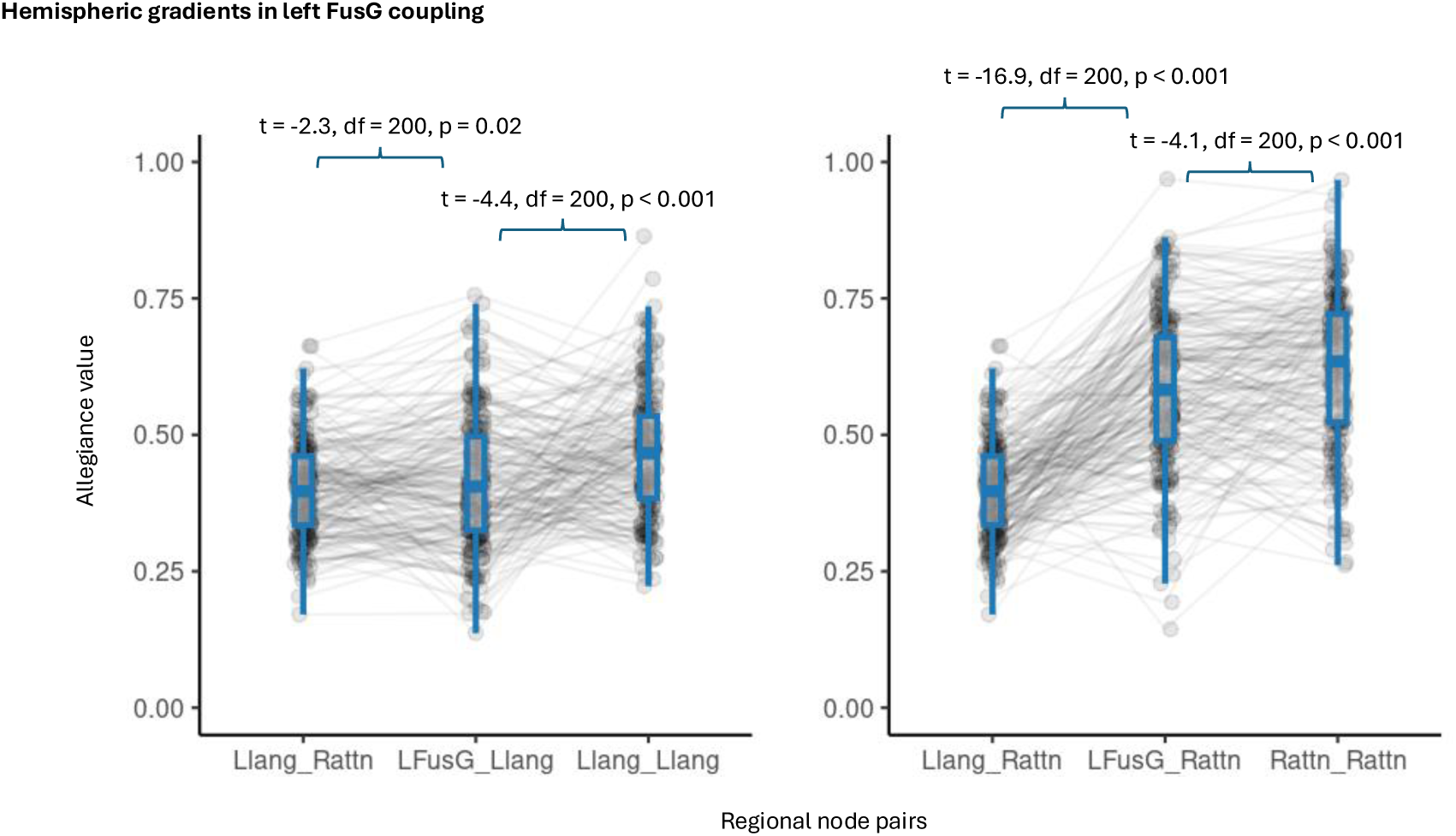
Hemispheric gradients in left FusG coupling (testing H2). Left: The lowest allegiance was observed between left language and right attention networks; intermediate allegiance between the left FusG and the left language network; and the highest allegiance within the left language network. Right: The lowest allegiance was again observed between left language and right attention networks; intermediate allegiance between the left FusG and the right attention network; and the highest allegiance within the right attention network.

### 3.3 Reading ability relates to greater left FusG-left IFG recruitment and reduced right FusG-left IFG integration

We next tested the hypothesis that reading ability would be positively associated with left FusG-left language recruitment, and negatively associated with right FusG-left language integration. We found no interaction effects between allegiance values and group, so we report results from the more parsimonious main-effects model. Regression coefficients with 95% confidence intervals are shown in Fig. 4b. Reading scores were associated with allegiance between the left FusG and left IFG ventral area 44, as well as between the right FusG and the same IFG node. Specifically, greater recruitment between the left FusG and left IFG ventral area 44 was associated with better reading performance, whereas greater integration between the right FusG and left IFG ventral area 44 was associated with poorer reading performance.

**Figure 4:**
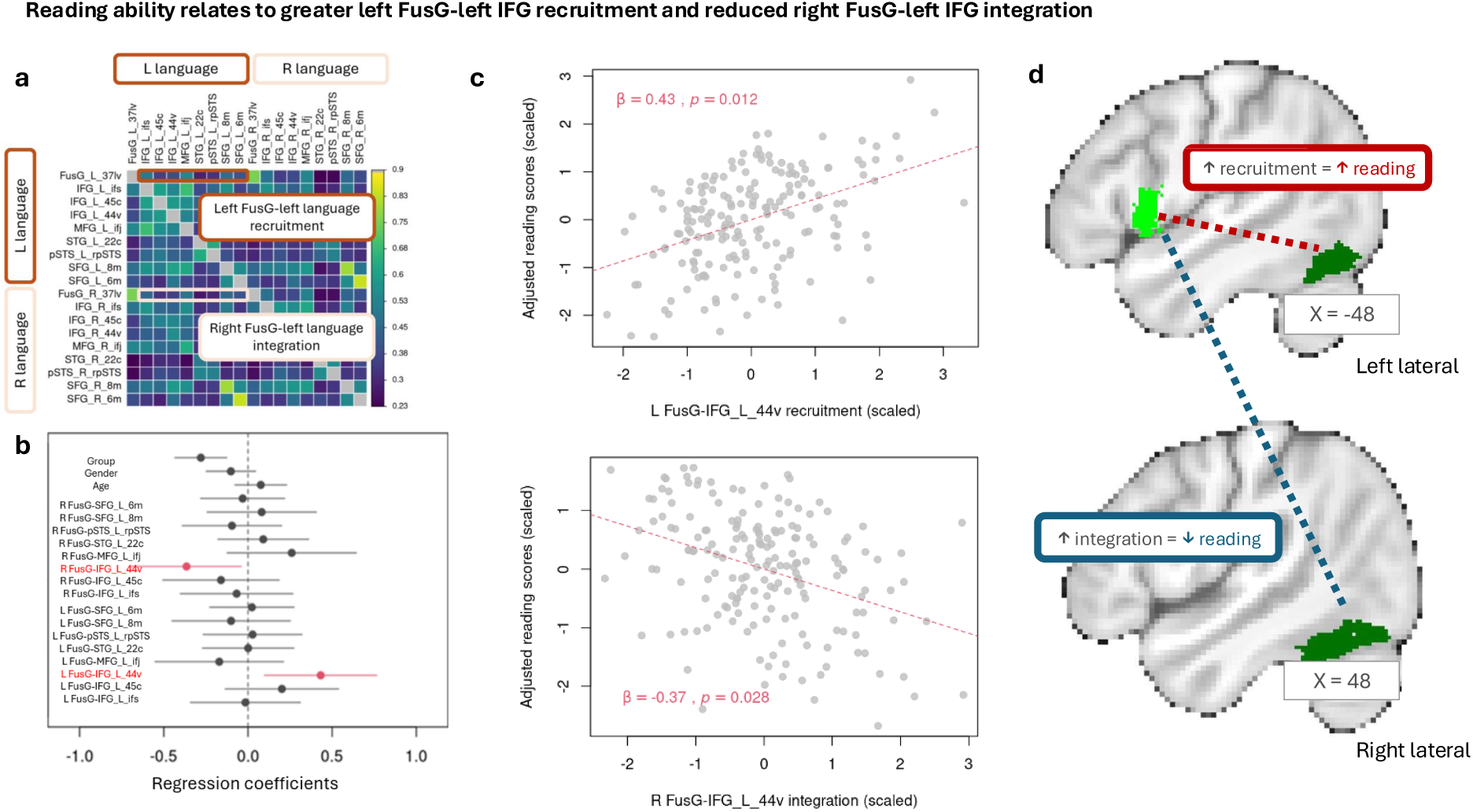
Reading is associated with greater recruitment of left FusG into the language system and weaker integration of right FusG into the language system (testing H3). a. Allegiance corresponding to the recruitment and integration of left and right FusG into the left language system. Diagonals all have value one and are shaded grey. b. Regression coefficients (dots) and their 95% confidence intervals (horizontal lines), indicating that recruitment of the left FusG into the left IFG ventral area 44 is positively associated with reading scores, whereas integration of the right FusG with the same region is negatively associated with reading scores. c. Adjusted reading scores plotted against left FusG-left IFG ventral area 44 recruitment (top). Adjusted reading scores plotted against right FusG-left IFG ventral area 44 integration (bottom). d. Schematic illustrations of the findings.

### 3.4 Effect of window lengths

The primary analysis was conducted using a window length of 40 TRs, as commonly used in previous studies. We repeated all analyses using window lengths of 20 and 60 TRs, and the corresponding results are presented in Supplementary Information Figures 1 and 2. The average module allegiance matrix obtained from the primary analysis (40-TR window) was highly correlated with those derived from the 20-TR window (Spearman’s *ρ* = 0.990, *p <* 0.001, two-sided) and the 60-TR window (Spearman’s *ρ* = 0.996, *p <* 0.001, two-sided), demonstrating consistency across these window lengths.

Despite this overall consistency, several differences are worth noting. First, when using a 20-TR window, left FusG-left language allegiance was significantly stronger than left language-right attention allegiance (H2; *p <* 0.001; Supplementary Information Fig. 1b). This difference did not survive thresholding in the primary analysis (*p* = 0.02; Fig. 3) or in the 60-TR analysis (*p* = 0.7; Supplementary Information Fig. 2b).

Second, in the reading score regression analysis (H3), the left FusG-left IFG ventral area 44 connection seen in the 40-TR analysis remained significant in the 60-TR analysis (*p* = 0.01; Supplementary Information Fig. 2c); while the same trend was present in the 20-TR analysis, this did not survive thresholding (*p* = 0.074; Supplementary Information Fig. 1c). The right FusG-left IFG ventral area 44 connection seen in the 40-TR analysis also showed a similar though non-significant trend in the 60-TR analysis (*p* = 0.06; Supplementary Information Fig. 1c), and was not significant in the 20-TR analysis (*p* = 0.40; Supplementary Information Fig. 1c). Overall, the pattern of results is broadly consistent across window lengths, though with hypothesised differences weakest using the 20-TR window, where fewer samples result in noisier connectivity estimates.

## 4 Discussion

In this study, we used MLNM to characterise the dynamic organisation of the human reading system in individuals with left-dominant language. Our three main findings were as follows. First, left language regions exhibited preferential intrahemispheric recruitment over interhemispheric integration, as also observed in Chai et al. (2016). We further compared this left intrahemispheric recruitment with that of their right hemisphere homologues, and found that left recruitment was indeed stronger. These findings indicate a distinctive left hemisphere organisation, which we suppose to relate to the specialised organisation for language in the left hemisphere. Second, we observed a graded allegiance pattern. Allegiance was lowest between left language and right attention regions, intermediate between the left FusG and each network, and highest within each network. This pattern supports the hypothesis that the left FusG flexibly couples with both left language and right attention nodes according to behavioural needs. Finally, reading ability was associated with the recruitment and integration of the FusG into the left language system. Specifically, better reading performance was linked to increased recruitment between the left FusG and left IFG, as well as reduced integration between the right FusG and left IFG. Together, these results highlight hemispheric asymmetries in the functional organisation of the reading system and provide novel evidence that individual differences in this organisation are related to reading ability.

### 4.1 Lateralisation of the reading networks

Our results converge with a large body of research demonstrating that reading is supported by a predominantly left-lateralised network (Price, 2012; Bonandrini et al., 2024). Using a MLNM approach, we extend this finding to the spatiotemporal domain, showing that left hemisphere language regions preferentially cluster together across time (Fig. 2b). This effect was unlikely to be driven by differences in task-related activation, as we regressed out the task component from each region’s time series.

Furthermore, we found that the left FusG preferentially forms modules with ipsilateral superior temporal regions relative to contralateral superior temporal regions, but it does not display a clear preference to organise with left or right inferior frontal regions. This pattern may reflect a sequential modular arrangement for decoding nonwords, whereby the left FusG initially couples with the left STG due to the STG’s role in phonological analysis (Price, 2012; Graves et al., 2008; Wise et al., 2001; Chang et al., 2010; Leonard & Chang, 2014; Skeide & Friederici, 2016). The left STG may then engage neighbouring regions and the left IFG to support the motor sequencing processes involved in pseudoword rhyming (Price, 2012; Sandak et al., 2004).

There are two observations that warrant further investigation. First, the left SFG medial area 6 does not show a preference for either intrahemispheric or interhemispheric coupling (Fig. 2b). In contrast, its neighbouring region, medial area 8, preferentially forms module with nearly all left language regions rather than integrating across hemispheres. Area 6 overlaps with the supplementary motor area and is primarily implicated in motor planning and initiation (Silva et al., 2018). Medial area 8 has been implicated in a range of functions, including the planning and coordination of eye movements (supplementary eye fields; Stuphorn, 2016), verbal working memory (Dadario et al., 2023), and resolution of uncertainty (Volzet al., 2004, 2005), all of which are potentially relevant to the paradigm employed. Both medial SFG regions have also been associated with maintaining cognitive set during task execution (Dosenbach et al., 2006; Alario et al., 2006; Crosson et al., 2018; Freedman et al., 1984). One possibility is that area 6 facilitates task processing by preparing both left language regions and their right homologues, whereas area 8 selectively modulates left-hemisphere language regions, such as through coordinating verbal working memory to resolve ambiguous rhymes, scanning back and forth between nonword pairs to support decisions. Second, the intrahemispheric recruitment between right IFG 44v and 45c is higher than that observed between their left hemisphere counterparts. A possible explanation is that the right IFG is involved in spatial attention and action execution (Hartwigsen et al., 2019), functions that are shared across both the pseudoword rhyming task and the pattern matching baseline.

### 4.2 FusG-frontal interactions and reading ability

The relationship between reading performance and FusG-IFG interactions highlights the previously reported importance of developmental specialisation of the left ventral occipitotemporal region in supporting reading ability. The left IFG ventral area 44 is strongly implicated in phonological processing, articulatory planning, and articulatory selection (Price, 2012; Sandak et al., 2004). These functions are critical for sounding out and comparing the phonological structure of pseudowords, as well as for supporting phonological working memory. This is especially relevant for tasks such as rhyme judgement, which require maintaining an auditory-phonological representation in working memory for comparison, a demand that is further heightened when processing pseudowords. Our data imply that stronger coupling between the orthographic processing system in FusG and the phonological processing system in ventral area 44 - i.e., tighter recruitment of the text analysis and speech planning systems into a common module - underpins proficient reading.

Furthermore, the observation that stronger left FusG-left IFG ventral area 44 recruitment (Fig. 4d) correlates with better reading is consistent with previous research showing that reading impairments in children are associated with reduced engagement of the left ventral occipitotemporal pathway and inferior frontal regions (Shaywitz et al., 2002). Conversely, the observation that stronger right FusG-left IFG integration is associated with poorer reading (Fig. 4d) suggests that persistent reliance on the right ventral occipitotemporal pathway may reflect an inefficient reading network. Although this effect was less robust across analyses using different window lengths (see Section 3.4), it remains noteworthy in light of converging evidence from the dyslexia literature, where compensatory right hemisphere activation has been linked to reduced reading efficiency in individuals with reading difficulties (Shaywitz et al., 2002; Tailby et al., 2014).

### 4.3 Left FusG as a multiplexing node

Using a psychophysiological interaction (PPI) analysis, we previously found that the left FusG up-regulates its connectivity with left language regions and right visuospatial attention regions depending on behavioural demands (He et al., 2025a). This pattern supports the view that the left FusG functions as a multiplexing node that flexibly couples with both left-hemisphere language and right-hemisphere attention systems (Chen et al., 2019). The present findings further reinforce this interpretation: the left FusG showed intermediate allegiance relative to the higher intranetwork allegiance observed within the left language and right attention core networks (implying flexible switching of FusG allegiance according to task), and to the lower internetwork allegiance between the left language and right attention networks (Fig. 3).

### 4.4 Methodological considerations

There are a number of methodological aspects that require further discussion. First, we used a whole-brain parcellation and included all nodes in the community detection to generate a reliable modular decomposition. Previous research in multilayer network modelling of language fMRI data has often focused exclusively on language regions while overlooking the FusG (e.g., Chai et al., 2016; He et al., 2018). A whole-brain approach provides broader coverage of network interactions, capturing regions that may indirectly support reading but are not traditionally classified as language regions. In addition, we used a 40 TRs (36 s) window for the main analysis of time-resolved connectivity and repeated all analyses with shorter (20 TRs, 18 s) and longer (60 TRs, 54 s) windows. The 40 TRs window aligns with prior recommendations that 30-60 s windows balance robustness and sensitivity (Hutchison et al., 2013; Shirer et al., 2012; Jones et al., 2012). Shorter windows are more susceptible to high frequency noise and longer ones are less sensitive to fast dynamics (Leonardi & Van De Ville, 2015; Zalesky & Breakspear, 2015). Finally, we used Pearson’s correlation to estimate the supra-adjacency matrix, as it is the most commonly used method (Pedersen et al., 2018a). Alternative approaches such as wavelet coherence (Bassett et al., 2013b; Chai et al., 2016; He et al., 2018), temporal-derivative methods (Shine et al., 2015, 2016), and phase synchrony analyses (Glerean et al., 2012), are also widely employed (Yang et al., 2021; Pedersen et al., 2018a).

### 4.5 Conclusion

Using multilayer network modelling of a language fMRI task, we revealed previously uncharacterised hemispheric asymmetries in an extended reading network and identified links between FusG-IFG interactions and individual reading ability. These results suggest that specific cross-regional dynamic coupling patterns may serve as neural markers of reading proficiency. Future longitudinal studies could investigate how these network configurations develop over time, providing insight into the neural mechanisms that support reading acquisition.

## Data and Code Availability

Any requests for access to the data used in this project should be directed to the Australian Epilepsy Project (a formal data access request can be lodged at https://www.epilepsyproject.org. au/research/access-to-aep-data). The multilayer community detection analysis was implemented using code from https://github.com/omidvarnia/Dynamic_brain_connectivity_analysis.

## Author Contributions

V.H.: Conceptualisation, Methodology, Formal Analysis, Writing—Original Draft, Writing—Review & Editing, and Visualisation. M.P.: Conceptualisation, Methodology, Writing-Review & Editing, and Funding Acquisition. D.N.V.: Conceptualisation, Methodology, Investigation, Resources, Writing-Review & Editing, Project Administration, and Funding Acquisition. H.R.P.: Investigation, Resources, Writing-Review & Editing, and Project Administration. J.E.C.: Investigation, Resources, Writing-Review & Editing, and Project Administration. G.D.J.: Conceptualisation, Methodology, Investigation, Resources, Writing-Review & Editing, Supervision, Project Administration, and Funding Acquisition. D.F.A.: Conceptualisation, Methodology, Software, Investigation, Resources, Data Curation, Writing-Original Draft, Writing-Review & Editing, Visualisation, Supervision, Project Administration, and Funding Acquisition. C.T.: Conceptualisation, Methodology, Investigation, Resources, Data Curation, Writing-Original Draft, Writing-Review & Editing, Visualisation, Supervision, Project Administration, and Funding Acquisition.

## Acknowledgments

The authors thank the Australian Epilepsy Project investigators (Supplementary Table 1). The Australian Epilepsy Project received funding from the Australian Government under the Medical Research Future Fund (Frontier Health and Medical Research Program - Grant Numbers MRFF75908 and RFRHPSI000008) and the Victoria State Government (Victorian-led Frontier Health and Medical Research Program). The Florey Institute of Neuroscience and Mental Health acknowledges the strong support from the Victorian Government and, in particular, the funding from the Operational Infrastructure Support Grant. The authors acknowledge the facilities and scientific and technical assistance of the National Imaging Facility, a National Collaborative Research Infrastructure Strategy (NCRIS) capability. This research was supported by The University of Melbourne’s Research Computing Services and the Petascale Campus Initiative. V.H. acknowledges the financial support received from the University of Melbourne through the Melbourne Research Scholarship. D.F.A. acknowledges fellowship funding from the National Imaging Facility.

