## SupplementaryInformation for "Multilayer Network Modelling of the Human Reading System"

### Supplementary Information:
### Multilayer Network Modelling of the Human Reading System

Vicky He[1,2], Mangor Pedersen[3], David N. Vaughan[1,2,4], Heath R. Pardoe[1,2], Jodie E. Chapman[1], Graeme D. Jackson[1,2,4], David F. Abbott[‡1,2,5], and Chris Tailby[‡1,2,6]

[1]The Florey Institute of Neuroscience and Mental Health, Heidelberg, Victoria, Australia

[2]Florey Department of Neuroscience and Mental Health, The University of Melbourne, Parkville, Victoria, Australia

[3]Department of Psychology and Neuroscience, Auckland University of Technology, Auckland, New Zealand

[4]Department of Neurology, Austin Health, Heidelberg, Victoria, Australia

[5]Department of Medicine-Austin Health, The University of Melbourne, Heidelberg, Victoria, Australia

[6]Department of Clinical Neuropsychology, Austin Health, Heidelberg, Victoria, Australia

**Corresponding authors:**

Vicky He

David F. Abbott

Chris Tailby

[‡]Joint senior authors

**List of Contents:**

Supplementary Figures 1-2

Supplementary Table 1

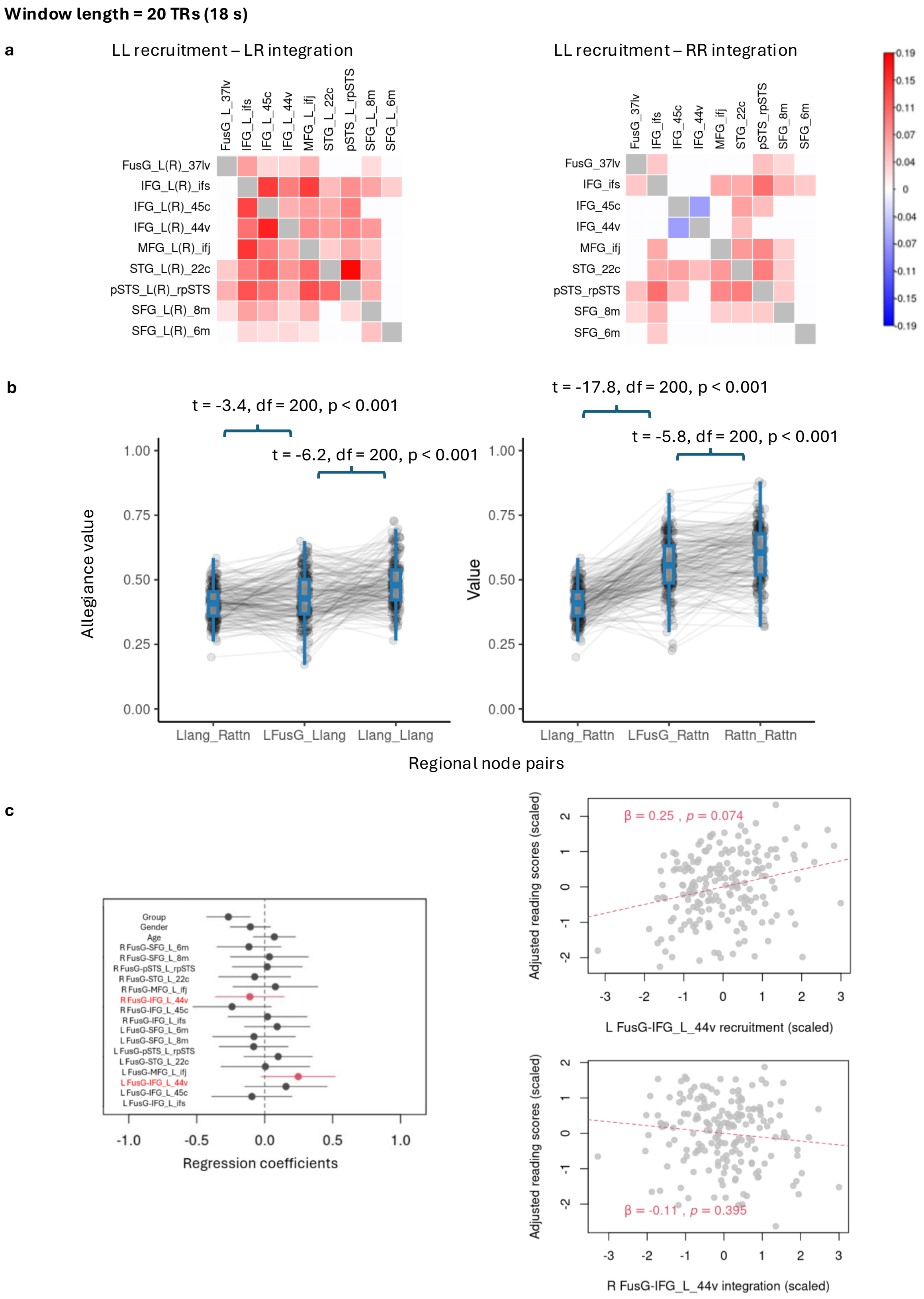


Figure 1: (Caption next page.)

Figure 1: Results obtained with window length 20 TRs (18 seconds). a. Left: Differences in probability between left language-left language recruitment and left language-right language integration. Right: Differences in probability between left language-left language recruitment and right language-right language recruitment. White cells denote non-significant results after Bonferroni correction. Diagonal cells are shaded grey, as they were not included in the analysis. b. Left: Hemispheric gradients in left FusG-left language coupling. Right: Hemispheric gradients in left FusG coupling-right attention coupling. c. Left: Regression coefficients (dots) and their 95% confidence intervals (horizontal lines). Right: Adjusted reading scores plotted against left FusG-left IFG ventral area 44 recruitment (top). Adjusted reading scores plotted against right FusG-left IFG ventral area 44 integration (bottom).

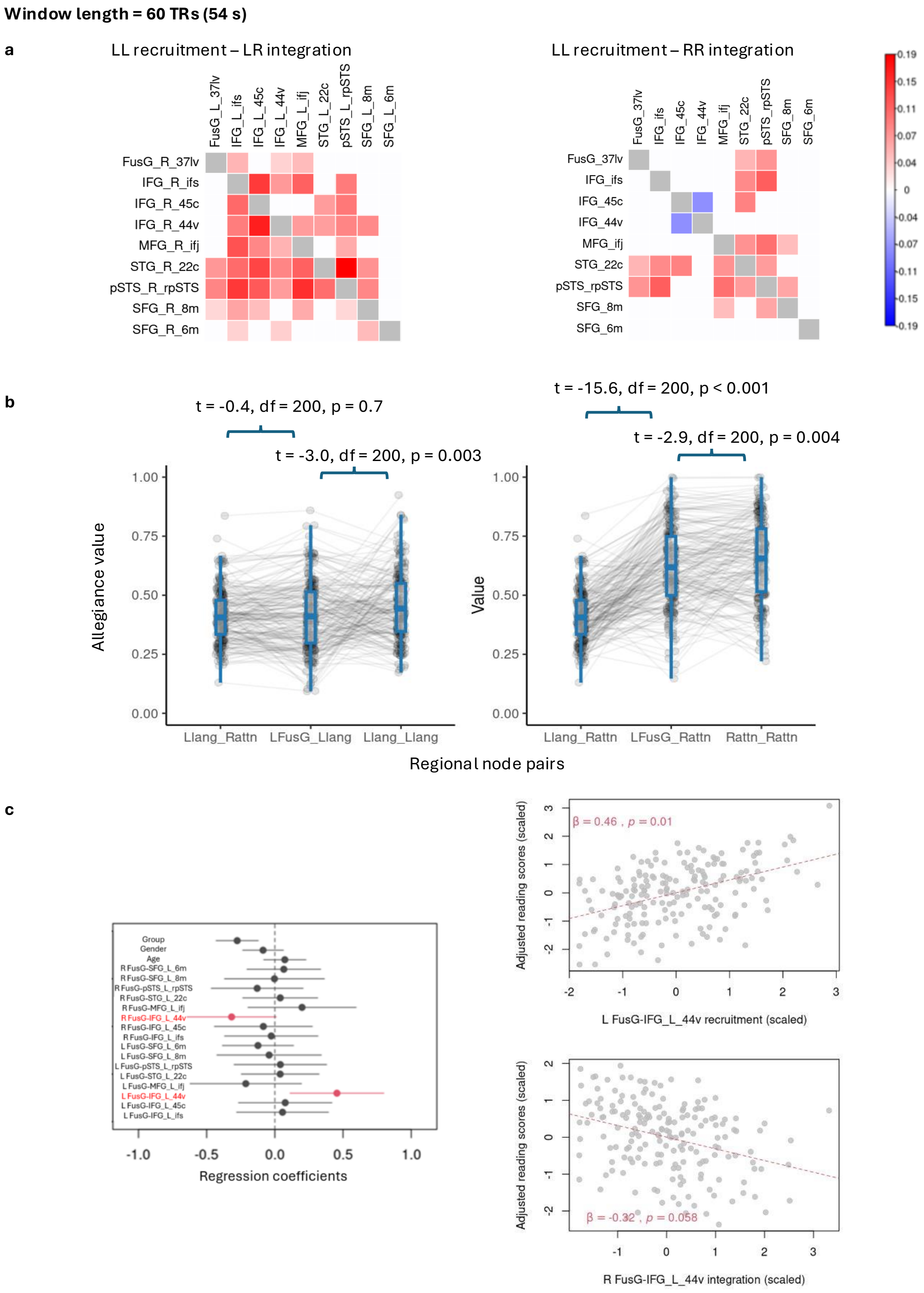


Figure 2: (Caption next page.)

Figure 2: Results obtained with window length 60 TRs (54 seconds). a. Left: Differences in probability between left language-left language recruitment and left language-right language integration. Right: Differences in probability between left language-left language recruitment and right language-right language recruitment. White cells denote non-significant results after Bonferroni correction. Diagonal cells are shaded grey, as they were not included in the analysis. b. Left: Hemispheric gradients in left FusG-left language coupling. Right: Hemispheric gradients in left FusG coupling-right attention coupling. c. Left: Regression coefficients (dots) and their 95% confidence intervals (horizontal lines). Right: Adjusted reading scores plotted against left FusG-left IFG ventral area 44 recruitment (top). Adjusted reading scores plotted against right FusG-left IFG ventral area 44 integration (bottom).

Table 1: Australian Epilepsy Project (AEP) investigator list with Contributor Roles Taxonomy (CRediT) author statements relevant for this manuscript.

| Name & ORCID | Primary Location | Role | CRediT Contribution |
|---|---|---|---|
| Graeme D. Jackson, MD 0000-0002-7917-5326 | The Florey Institute of Neuroscience and Mental Health | Chief Investigator | Conceptualisation; Methodology; Investigation; Resources; Writing – Review & Editing; Supervision; Project Administration; Funding Acquisition |
| David F. Abbott, PhD | | | |

*Continued on next page*

*Continued from previous page*

| **Name & ORCID** | **Primary Location** | **Role** | **CRediT Contribution** |
|---|---|---|---|
| 0000-0002-7259-8238 | The Florey Institute of Neuroscience and Mental Health | Informatics Lead | Conceptualisation; Methodology; Software; Investigation; Resources; Data Curation; Writing – Original Draft; Writing – Review & Editing; Visualisation; Supervision; Project Administration; Funding Acquisition |
| Zanfina Ademi, PhD | | | |
| 0000-0002-0625-3522 | Monash University | Health Economics Lead | Conceptualisation; Funding Acquisition |
| Subhaga Amarasekara | | | |
| – | The Florey Institute of Neuroscience and Mental Health | Product Lead | Resources; Project Administration |
| Amanda Anderson | | | |

*Continued on next page*

*Continued from previous page*

| **Name & ORCID** | **Primary Location** | **Role** | **CRediT Contribution** |
|---|---|---|---|
| – | The Florey Institute of Neuroscience and Mental Health | Lived Experience Ambassador and Participant Lead | Investigation; Resources; Funding Acquisition |
| Rachel Hughes<br>– | The Florey Institute of Neuroscience and Mental Health | Clinical Research Coordinator | Investigation; Resources |
| Donna Hutchison<br>– | The Florey Institute of Neuroscience and Mental Health | Executive Lead | Project Administration |
| Patrick Kwan, MD<br>0000-0001-7310-276X | Monash University | Outcomes Lead | Conceptualisation; Resources; Funding Acquisition |
| Paul Lightfoot<br>– | The Florey Institute of Neuroscience and Mental Health | Operations Lead | Investigation; Project Administration |

*Continued on next page*

*Continued from previous page*

| **Name & ORCID** | **Primary Location** | **Role** | **CRediT Contribution** |
|---|---|---|---|
| Saul Mullen, MD, PhD<br>0000-0003-1224-4101 | The University of Melbourne | Protocol Development Lead (2019–2021) | Conceptualisation; Methodology; Funding Acquisition |
| Karen L. Oliver, PhD<br>0000-0001-5188-6153 | The University of Melbourne | Genetics Lead | Conceptualisation; Funding Acquisition |
| Heath R. Pardoe, PhD<br>0000-0002-0123-2167 | The Florey Institute of Neuroscience and Mental Health | Science Operations Lead | Investigation; Resources; Writing – Review & Editing; Project Administration |
| Mangor Pedersen, PhD<br>0000-0002-9199-1916 | Auckland University of Technology | Artificial Intelligence Lead | Conceptualisation; Methodology; Funding Acquisition |

*Continued on next page*

*Continued from previous page*

| **Name & ORCID** | **Primary Location** | **Role** | **CRediT Contribution** |
|---|---|---|---|
| Chris Tailby, PhD 0000-0002-1320-5924 | The Florey Institute of Neuroscience and Mental Health | Neuropsychology Lead | Conceptualisation; Methodology; Investigation; Resources; Data Curation; Writing – Original Draft; Writing – Review & Editing; Visualisation; Supervision; Project Administration; Funding Acquisition |
| David N. Vaughan, MD, PhD 0000-0002-6225-7739 | The Florey Institute of Neuroscience and Mental Health | Imaging Lead | Conceptualisation; Methodology; Investigation; Resources; Writing – Review & Editing; Project Administration; Funding Acquisition |
| Anton De Weger | | | |

*Continued on next page*

*Continued from previous page*

| Name & ORCID | Primary Location | Role | CRediT Contribution |
|---|---|---|---|
| 0009-0006-7478-361X | The Florey Institute of Neuroscience and Mental Health | Digital and Technology Lead | Software; Resources; Data Curation |
